# Impaired effort perception, not reward sensitivity, impacts the effort-reward trade-off in Parkinson’s disease

**DOI:** 10.64898/2026.03.26.714286

**Authors:** Jonathan M. Wood, Alyssa N. Eyssalenne, Amanda S. Therrien, Aaron L. Wong

## Abstract

Deciding whether and how to act depends on a trade-off between the effort required to execute a given action and the potential reward for completing it. Impairments in this effort-reward trade-off have been proposed to underlie reduced movement vigor, or bradykinesia, in Parkinson’s disease (PD). However, several mechanisms could alter the effort-reward trade-off in PD, each with unique implications for understanding and treating bradykinesia. Therefore, we individually examined whether people with PD (both on and off dopamine medication) demonstrated reduced sensitivity to reward value, increased perception of effort, or a biased mapping between effort and reward, compared to age- and sex-matched neurotypical controls. We found that people with PD exhibited increased effort perception and, surprisingly, no reduced sensitivity to reward value or a biased mapping between effort and reward. These findings suggest that effort perception could be an important factor driving bradykinesia in PD.

## Background

Deciding whether and how to act depends, at least in part, on a trade-off between the effort required for a given task and the potential reward for completing it. This is exemplified by the stark contrast between a child eagerly leaping out of bed on Christmas morning and their reluctance to rise before a big test at school. A large body of literature shows that this effort-reward trade-off is critical for motivating our actions: we are willing to make more vigorous movements – i.e., move faster^1–5^ or more forcefully ^6,7^, make quicker decisions^8–10^, and perform more difficult cognitive tasks^11^ – given the prospect of greater reward.

Changes to the effort-reward trade-off have been proposed to underlie the reduced movement vigor, termed bradykinesia, characteristic of Parkinson’s disease (PD)^12–14^. This is based on the finding that people with PD retain the capacity to produce greater forces and faster movement speeds when sufficiently motivated^14,15^. For a given magnitude of reward, however, people with PD are less willing to produce the same amount of force as neurotypical controls^16–22^. Despite evidence of an impaired effort-reward trade-off in PD that could affect movement vigor, the underlying cause of this impairment is unclear. Critically, there are several potential mechanisms through which PD could alter the effort-reward trade-off, each with unique implications for understanding and potentially treating bradykinesia.

An impaired effort-reward trade-off in PD could stem from reduced sensitivity to the rewards offered in exchange for effort production^18,19^. From this standpoint, bradykinesia could be remedied simply by increasing movement incentives. PD has long been thought to reduce reward sensitivity, likely because of the overlap between its pathophysiology – degeneration of dopamine-producing cells in the substantia nigra^23–25^ – and the role of midbrain dopamine circuits in reward processing^26–28^. In support of this view, people with PD have difficulty learning from reward feedback^29–31^. Yet, an effort-reward trade-off hinges on established (i.e., baseline) sensitivities to the value of potential movement outcomes rather than on learning novel reward values^16–22^. Because prior studies used monetary rewards to motivate effortful actions, we operationalize reward sensitivity as an individual’s subjective value of a given monetary reward. Many studies have used a decision-making task^32,33^ to assess sensitivity to different monetary reward values in people with PD^34–37^. Yet these studies have typically provided individuals with a continuous tally of their “earnings,” affording the opportunity to learn from previous decisions. Thus, any differences in reward sensitivity between PD and controls may be confounded by differences in the ability to learn from reward outcomes. Whether PD specifically alters reward sensitivity and how this may impact the effort-reward trade-off in this population remains unclear.

A reduction in reward sensitivity is not the only mechanism by which PD could alter the effort-reward trade-off. PD could also cause movements to be perceived as more effortful than they truly are, thus requiring greater rewards to compensate. In this case, bradykinesia would reflect a mismatch between actual and perceived effort, such that an individual may perceive they are exerting more force than they truly are. Effort perception is typically tested in one of two ways. The first has participants match a prescribed force level, treating the sense of effort as a product of peripheral somatosensation and the central processing of ongoing motor commands^38–42^. The other method has participants retrospectively rate perceived effort following force production^43–46^, effectively assessing cognitive beliefs about one’s effort. Interestingly, peripheral deafferentation does not impair force-matching performance despite altering cognitive beliefs about perceived exertion^47^, suggesting that these two measures of effort might reflect distinct perceptual phenomena. While PD is known to impair somatosensation^48–50^, the impact on the perception of effort and whether it differentially affects the sensory processing of motor commands and cognitive beliefs about exertion remains unknown.

Finally, PD could selectively distort the mapping between effort and reward. That is, rather than directly changing reward sensitivity or effort perception, bradykinesia in PD could reflect a bias in the internal calibration between these two components. A biased mapping would appear as a systematic difference in the rate at which people with PD are willing to increase effort production in response to rewards of increasing magnitude (i.e., the slope of the effort-reward relationship), compared to neurotypical controls. Prior work examining the effort-reward mapping has yielded mixed results^16,17,22^, perhaps because participants were required to produce forces throughout the experimental task and may have experienced muscular fatigue. Fatigue can induce a paradoxical increase in perceived exertion despite decreases in actual force production^39^, creating the appearance of a distorted effort-reward mapping, especially at higher effort levels. Hence, it remains unclear exactly how PD biases the mapping between effort and reward.

Here, we systematically investigated the factors that could underlie an altered effort-reward trade-off in PD by comparing reward sensitivity, effort perception (assessed using both sensorimotor- and cognition-based measures), and the slope of the effort-reward relationship between individuals with PD and neurotypical control participants. Because dopamine has been implicated in reward sensitivity, we additionally compared each factor within the PD group across two conditions: on and off their dopaminergic medication. Our results showed that PD impairs effort perception, such that exerted forces were perceived as greater during both force matching and retrospective ratings of perceived exertion. In contrast, reward sensitivities and effort-reward relationship slopes were similar in the PD and control groups, regardless of medication status. These results add important context to prior findings of distorted effort-reward trade-offs in people with PD, and have critical implications for understanding and treating bradykinesia in this population.

## Results

The goal of this study was to examine three potential factors contributing to effort-reward trade-off impairments in people with PD: 1) a reduced sensitivity to rewards, 2) an increased perception of effort, or 3) a biased effort-reward mapping. To test these factors, participants with PD (N=37) and neurotypical controls (N = 39) performed three separate tasks using a Kinarm Exoskeleton robot (BKIN Technologies, ON, Canada). Thirty-two participants with PD were tested both soon after taking their dopaminergic medication (the ‘ON’ medication state) and after skipping a single medication dose (the ‘OFF’ medication state). Of the remaining participants in the PD group, two completed testing only in the ON state, and three completed testing only in the OFF state. People in the ON medication state took their last dose 2.0 ± 1.3 hours (mean ± 1 SD) prior to testing; people in the OFF medication state took their last dose 14.7 ± 3.8 hours before testing. Removing participants, n = 7, who completed OFF testing <12 hours after their last dose did not change the results. A detailed breakdown of PD group demographics, including the precise time since last medication dose for the ON and OFF testing conditions, is shown in the Supplementary Table 1. Our sample of participants with PD was in the early stage of the disease process as measured by the Hoen and Yahr score (2.0 ± 0.5). Disease severity was quantified using the United Parkinson’s Disease Rating Scale motor score (UPDRS Part III = 32.2 ± 13.5) which showed no reliable relationship with any of our primary outcome measures. As a group, our sample of people with PD was mildly bradykinetic as determined by the speed of their reaching movements relative to controls (see Supplemental Results; group difference in reach velocity normalized by reach amplitude, PD ON vs controls mean [95% HDI] = −0.42 [−0.71 −0.12], p_difference_ = 99.7%, d = −0.69; PD OFF vs controls = −0.59 [−0.88 −0.29], p_difference_ = 100.0%, d = −0.96). People with PD also moved more slowly when OFF their dopamine replacement medication compared to ON (−0.18 [−0.28 −0.07], p_difference_ = 99.8%, d_z_ = −0.70).

All statistical analyses were conducted using Bayesian t-tests (see Methods) to examine differences in group means^51,52^. The size of each group-mean difference was quantified as the mean and 95% high-density interval (HDI) – i.e., the interval containing the true difference with 95% certainty. We also quantified the reliability of this group difference (p_difference_) as the percentage of the posterior distribution falling on one side of 0; higher p_difference_ values indicate a greater probability of a true difference between group means. We considered both the size and reliability of the effect together when interpreting our results, with p_difference_ values >95% being considered sufficiently reliable. For each comparison, we calculated a posterior distribution of standardized effect size (Cohen’s d or d_z_ for between or within subjects comparisons, respectively) and report the mean of that distribution. For each comparison, we were primarily interested in three effects: the effect of disease with medication (a between-subjects comparison of PD ON and controls), the effect of disease without medication (a between-subjects comparison of PD OFF and controls), and the effect of medication state (a within-subjects comparison of PD ON and PD OFF). All data and code are openly available at https://osf.io/krw49.

### Reward sensitivity was similar between people with PD and neurotypical controls

We first tested the hypothesis that PD decreases reward sensitivity using a conventional decision-making task from the behavioral economics literature (see Methods)^32,33,53,54^. Participants indicated their preference between two options (Fig. 1a): one in which winning a monetary reward was guaranteed (i.e., the certain option) and another in which there was a 50/50 chance of winning a monetary reward or winning nothing (i.e., the gamble option). Prior to beginning the experiment, participants were informed of the gamble probability and that a portion of their winnings would be added to their total compensation for study participation. Importantly, to mitigate the confounding effect of learning from their choices, participants were not informed about the outcome of chosen gambles, their current cumulative reward, or the number of trials left in the task.

**Fig. 1:**
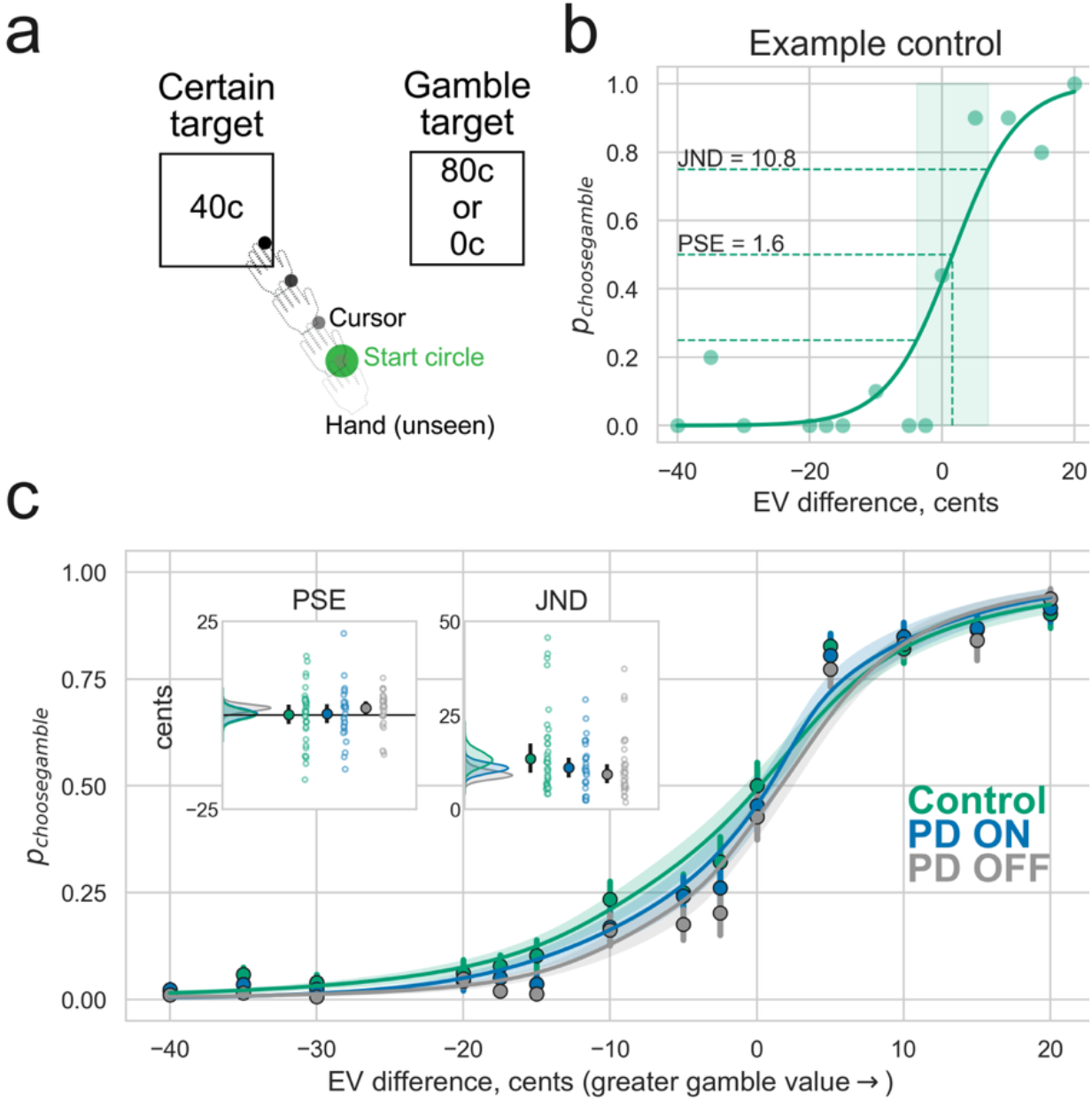
Reward sensitivity. **(a)** Participants made decisions by reaching to either a certain target (guaranteed win) or a gamble target (50/50 chance of winning a monetary reward or nothing). We calculated expected value (EV) differences for each choice (Gamble – Certain). **(b)** Example control participant that was closest to the median of all PSEs. Empirical probability of choosing the gamble at each EV level (dots) along with the fitted logistic function (solid line). **(c)** The empirical data averaged across the participants and groups (circles) against the mean logistic functions created from individual parameter values. Error bars and shading represent 1 SEM. For this analysis, we removed 12 participants (4 controls, 8 PD: 3 ON only, 3 OFF only and 2 both ON and OFF) because their data could not be fit with a logistic function (see Methods). **(insets)** Kernel density estimates of the posterior means for each group’s PSE and JND, plotted alongside the empirical group means (filled circles) and individual datapoints (open circles). Error bars are the 95% HDIs from each posterior distribution.

To quantify reward sensitivity, we computed the point of subjective equality (PSE) on a psychometric function fit to participants’ choice data (see Methods). The PSE represents the inflection point where participants switched their preference from the certain to the gamble option. A positive PSE indicated that participants preferentially chose the gamble option when its expected value was greater than the certain option, while a negative PSE indicated the opposite. Hence, more positive PSEs reflected more conservative behavior, consistent with reduced reward sensitivity. In contrast, more negative PSEs were associated with choosing the gamble option more often, consistent with increased sensitivity to potential rewards.

We found that the PSE values were similar between the groups (Fig. 1c). The PSE differences were extremely small and not reliable between PD ON and controls (mean difference [95% HDI] = 0.3 cents [−3.3 3.9], p_difference_ = 55.4%, d = 0.04), PD OFF and controls (1.7 cents [−1.4 4.9], p_difference_ = 86.0%, d = 0.29), and PD ON and PD OFF (0.6 cents [−1.01 2.3], p_difference_ = 80.1%, d_z_ = 0.21). Because of this surprising null result, we examined the strength of evidence supporting the null hypothesis using a Bayes factor (BF) analysis^55,56^. We found moderate evidence for equal PSE values between PD ON and controls (BF_10_ = 0.26) and between PD ON and PD OFF (BF_10_ = 0.27), but weaker evidence for the PD OFF vs control comparison (BF_10_ = 0.39). We also observed no difference in the degree to which movement vigor (i.e., reaching speed) changed across different magnitudes of reward on offer (Supplemental Figure S1), consistent with the idea that people with PD did not respond differently to reward. Therefore, contrary to theories about PD and reward processing, we did not find evidence that PD and reduced dopamine availability decreased reward sensitivity.

Decreased reward sensitivity could also manifest as greater noise in decision making between the certain and gamble options. Thus, as a secondary analysis, we calculated the just noticeable difference (JND) as the change in the expected value difference between the 75^th^ and 25^th^ percentiles on the psychometric function (Fig. 1c right inset). If people with PD were less sensitive to changes in reward value, they should exhibit larger JNDs on average. However, this was not the case. The difference in JND values was small and unreliable between PD ON and controls (−2.4 cents [−6.9 2.2], p_difference_ = 85.3%, d = −0.30), between PD OFF and controls (−2.6 cents [−6.5 1.1], p_difference_ = 92.0%, d = −0.43) and between PD ON and PD OFF (0.3 cents [−2.5 3.0], p_difference_ = 59.0%, d_z_ = 0.06). Thus, we did not observe an impact of PD or dopamine availability on the sensitivity to changes in reward value.

When fitting a psychometric function to choice data in the reward sensitivity task, not all participants met goodness-of-fit criteria (n=12, see Methods). To rule out the possibility that their exclusion drove the results above, we performed an additional analysis that allowed us to include all participants. Specifically, we calculated the probability of choosing the gamble option when the gamble and fixed options had the same expected value (EV difference = 0). Here, a lower probability of selecting the gamble option indicated lower reward sensitivity. These results aligned with our primary analysis: the differences in the probability of choosing the gamble option were small and not reliable (PD ON vs control = −0.07 [−0.23 0.09], p_difference_ = 79.8%, d = −0.20; PD OFF vs control = −0.08 [−0.23 0.07], p_difference_ = 86.3%, d = −0.26; PD ON vs PD OFF = 0.02 [−0.06 0.09], p_difference_ = 65.7%, d_z_ = 0.08). Using the Bayes Factor analysis, we found moderate evidence for equality between PD ON and PD OFF (BF_10_ = 0.23), but the evidence for the null was weaker for PD ON vs controls (BF_10_ = 0.33) and for PD OFF (BF_10_ = 0.40). Taken together, these analyses do not support the hypothesis that reward sensitivity is impaired in people with PD or that it is influenced by dopamine availability.

### The sensorimotor perception of effort was greater in people with PD

Second, we tested the hypothesis that PD influences the perception of effort. Using a force-matching task in which participants produced isometric elbow extension forces, we assessed the sensorimotor-based perception of effort, thought to arise from a combination of peripheral and central processing mechanisms^38–42^ (see Methods). We chose an extension force because PD typically affects the production of extension movements more than flexion movements^57^. During each trial (Fig. 2a), participants first produced an elbow extension force guided by visual feedback (the initial force; Fig. 2b). After a brief rest, they were asked to reproduce the same force with the same arm without visual feedback (the match force; Fig. 2c). Since there was no visual feedback to guide the match force, we assume that participants perceived the effort produced during the match force to be equal to that of the initial force. A lower match force, therefore, indicated that participants perceived the effort associated with the match force to be greater than its true magnitude (i.e., an increased sensorimotor perception of effort).

**Fig. 2:**
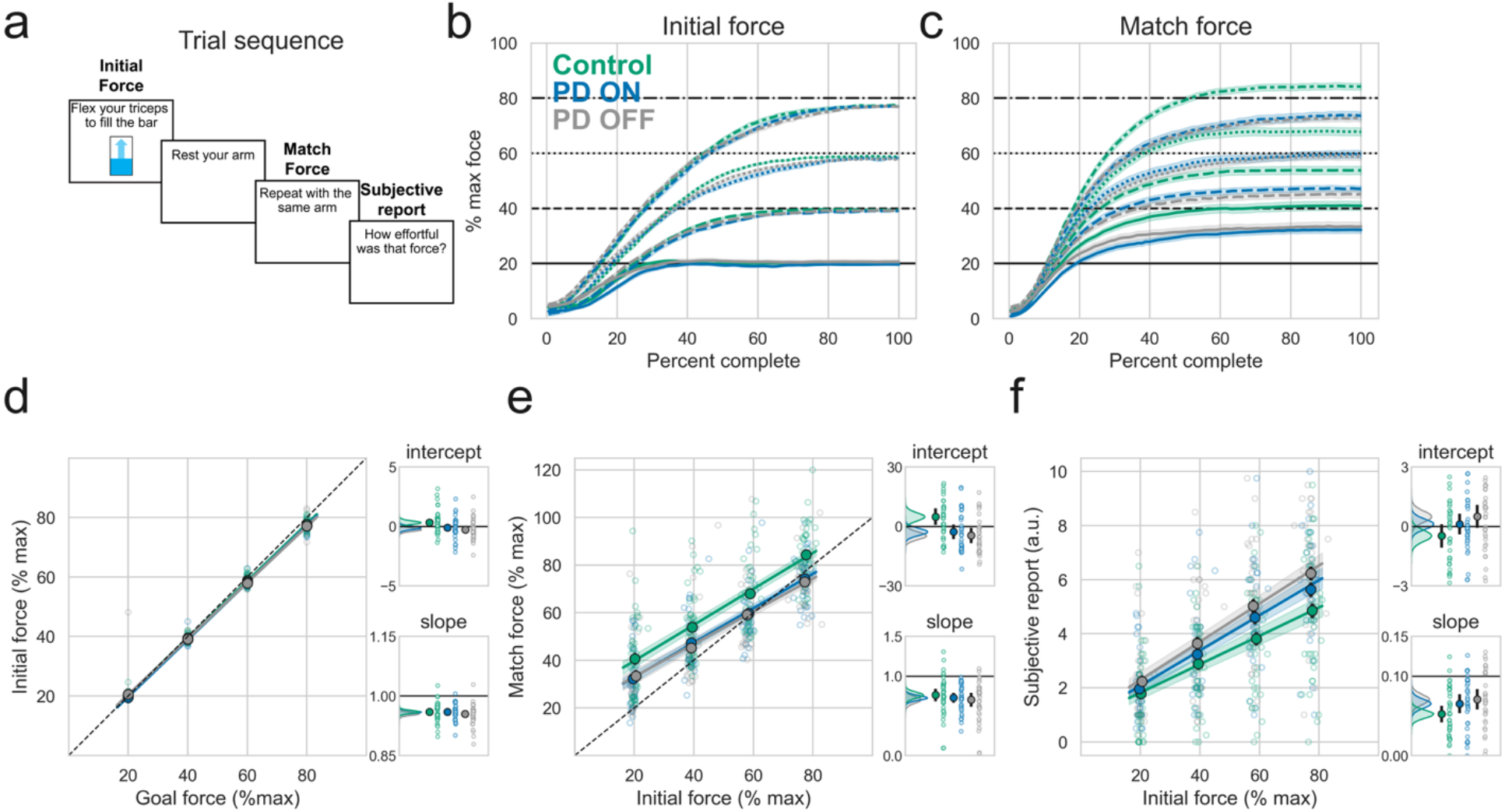
Perception of effort. **(a)** Task sequence. Participants first produced an initial force guided by visual feedback to one of 4 different force levels (20, 40, 60, or 80% max force). After a brief rest, they were asked to produce the same force with the same arm, this time without any visual feedback. After resting, they were asked to rate how effortful the movement was on a scale of 0 to 10 (10 being their max effort and 0 being no effort). Mean initial **(b)** and match **(c)** force traces for each group (colors) and each goal force, denoted with different line styles. For the purposes of these plots, each participant’s force trace data was averaged at each percentile so everyone had 100 data points. Shading represents the standard error of the mean (SEM), and horizontal black lines represent each goal force, with a different style line for each goal. **(d-f, main panels)** Each outcome of interest (y-axis) is plotted against the goal or initial forces. Solid circles with black outlines represent the overall group means, error bars (not visible for all datapoints) represent 1 SEM, and open circles represent individual participant means at each goal force level. The solid lines are the mean regression lines for each group, and shading represents 1 SEM. **(d-f, inset panels)** Kernel density estimates of the posterior distribution of group means for the intercepts (top) and slopes (bottom), calculated using Bayesian t-tests. The empirical group means (filled circles) and individual datapoints (open circles) are plotted to the right. Error bars for these plots (not visible for all datapoints) reflect the 95% HDIs for each respective posterior distribution. Note that the intercepts are calculated from the mean-centered data. **(d)** Performance on the initial forces, when the feedback was visible, was similar across groups and conditions. **(e)** Match forces were lower in the PD ON group than in controls, indicating an increase in the sensorimotor-based perception of effort. **(f)** Subjective effort ratings were higher and increased faster for the PD OFF group compared to controls (Supplemental Results).

We first confirmed that performance on the initial force, when the feedback was visible, was similar between groups (Fig. 2d). We regressed each participant’s mean-centered initial forces against the respective mean-centered goal forces. The resulting intercepts reflected the initial force (relative to the mean) produced at the average goal force level, and the slope reflected the change in the initial force across the goal force levels. The groups did not show meaningful differences in initial force production (Fig. 2b & d). While two of the intercept differences were reliable, we did not interpret them as meaningful given their small sizes (PD ON vs controls = −0.5% max force [−0.9 −0.04], p_difference_ = 98.4%, d = −0.56; PD OFF vs controls = −0.6% max force [−0.9 −0.2], p_difference_ = 99.6%, d = −0.8; PD ON vs PD OFF = 0.1% max force [−0.3 0.4], p_difference_ = 69.1%, d_z_ = 0.10). Slope differences were not large or reliable (PD ON vs controls = −0.001 [−0.01 0.01], p_difference_ = 54.9%, d = −0.03; PD OFF vs controls = −0.01 [−0.02 0.005], p_difference_ = 86.8%; d = −0.33; PD ON vs PD OFF = 0.01 [−0.005 0.01], p_difference_ = 84.3%, d_z_ = 0.19). Therefore, the performance of initial forces, when the feedback was visible, was similar across groups and conditions.

We next analyzed the match forces, our primary measure of the sensorimotor-based perception of effort. We regressed the mean-centered match force data against the mean-centered initial force data for each participant. Here, the intercept reflected the match force (relative to the mean) produced at the average initial force level. Comparison across groups showed that the PD ON group exhibited much lower intercepts than the control group (Fig. 2e; −7.6% max force [−13.0 −2.0], p_difference_ = 99.6%, d = −0.66), as did the PD OFF group compared to controls (−9.1% max force [−14.6 −3.6], p_difference_ = 99.9%, d = −0.83). While the PD ON condition had numerically greater intercepts compared to PD OFF, this difference was not reliable (2.5% max force [−0.8 5.9], p_difference_ = 92.8%, d_z_ = 0.28). To assess whether this increased effort perception was consistent across force levels, we examined the difference in regression slopes. Slope differences were not reliable for any of the three comparisons (PD ON vs controls = −0.04 [−0.13 0.07] p_difference_ = 75.1%, d = −0.17; PD OFF vs controls = −0.06 [−0.18 0.06], p_difference_ = 85.3, d = −0.26; PD ON vs PD OFF = 0.05 [−0.01 0.10], p_difference_ = 93.7%, d_z_ = 0.29). Together, these results suggested that people with PD showed an increase in the sensorimotor-based perception of effort, which was not significantly impacted by dopamine availability.

It was important to test whether physical fatigue may have contributed to a decrease in match forces in the PD group, especially for the PD ON and control comparison. In our task, physical fatigue could manifest as a decrease in initial forces and/or match forces across trials. To assess the former, we calculated the difference between each initial force and each respective goal force, then regressed this difference against trial number for each participant. To assess match forces, we performed the same regression using the difference between the match forces and each respective goal force. For both analyses, more negative slopes would indicate a progressive decrease in force output across trials, suggesting greater fatigue. For the initial forces, we did not observe differences in the slopes between the PD ON and control groups (0.004 [−0.03 0.04], p_difference_ = 58.1%, d = 0.06) or between the PD ON and PD OFF conditions (0.02 [−0.02 0.06], p_difference_ = 82.8%, d_z_ = 0.22). While the PD OFF group did reliably decrease their initial forces across trials more than controls (−0.04 [−0.07 0.003], p_difference_ = 96.3%, d = −0.57), this small effect cannot fully explain the large differences in match force intercepts between the PD and control groups. We found similar results for the match force analysis (PD ON vs controls = 0.14 [−0.35 0.60], p_difference_ = 73.5%, d = 0.18; PD ON vs PD OFF = −0.10 [−0.51 0.30], p_difference_ = 69.9%, d_z_ = −0.11; PD OFF vs controls = 0.37 [−0.03 0.74], p_difference_ = 96.9%, d = 0.58). Overall, these results indicate that fatigue, as measured by progressive changes in either the initial or match forces, could not account for the differences in sensorimotor effort observed above.

### The cognitive perception of effort was increased in people with PD

Cognitive perception of effort was measured by asking participants to retrospectively rate the level of effort they perceived to have exerted after each trial in the force-matching task. We analyzed the subjective effort reports by regressing each participant’s mean-centered subjective effort reports against the respective mean-centered initial forces (Fig. 2f). Thus, the intercept reflected the subjective effort report (relative to the mean) at the average force level, and the slope reflected the change in the cognitive perception of effort as the goal force increased. We observed greater intercepts and slopes for the PD ON group compared to controls, although these differences were not reliable (intercept difference = 0.6 [−0.2 1.4], p_difference_ = 92.9%, d = 0.36; slope difference = 0.01 [−0.003 0.03], p_difference_ = 94.7%, d = 0.40). Likewise, the intercepts and slopes were lower for the PD ON condition compared to PD OFF, but not reliably so (intercept difference = −0.5 [−1.0 0.2], p_difference_ = 93.8%, d_z_ = −0.30; slope difference = −0.01 [−0.02 0.01], p_difference_ = 87.1%, d_z_ = −0.21). Both the intercept and slope were reliably greater for the PD OFF group relative to controls (intercept difference = 1.0 [0.2 1.8], p_difference_ = 98.9%, d = 0.56; slope difference = 0.02 [0.002 0.04], p_difference_ = 98.5%, d = 0.53). Combined, these results suggest that having PD and being off dopamine medication together increase the cognitive perception of effort.

### The mapping between effort and reward was similar between people with PD and neurotypical controls

Lastly, we tested the hypothesis that PD selectively biases the mapping between effort and reward. To assess this mapping, participants performed a decision-making task (Fig. 3a) in which they were asked if they would be willing to produce a certain amount of effort (an isometric elbow extension force) in exchange for a monetary reward. As fatigue can alter the willingness to exert effort over time^58,59^, participants were not required to immediately perform accepted trials; instead, 5 of the accepted trials were randomly selected and performed at the end of the task. Across all force and reward levels, the PD OFF group was less likely to accept trials compared to controls (−0.05 [−0.11 0.01], p_difference_ = 95.0%, d = −0.41) although we consider this a small effect given it equates to about 6 fewer accepted trials in this task. The difference between the PD ON group and controls was not reliable (−0.04 [−0.10 0.02], p_difference_ = 91.3%, d = −0.34). There were no differences in acceptance rate between PD ON and PD OFF conditions (0.01 [−0.05 0.07], p_difference_ = 61.9%, d_z_ = 0.06). In general, there was little difference in overall willingness across groups to accept performing effort in exchange for reward in this task.

**Fig. 3.**
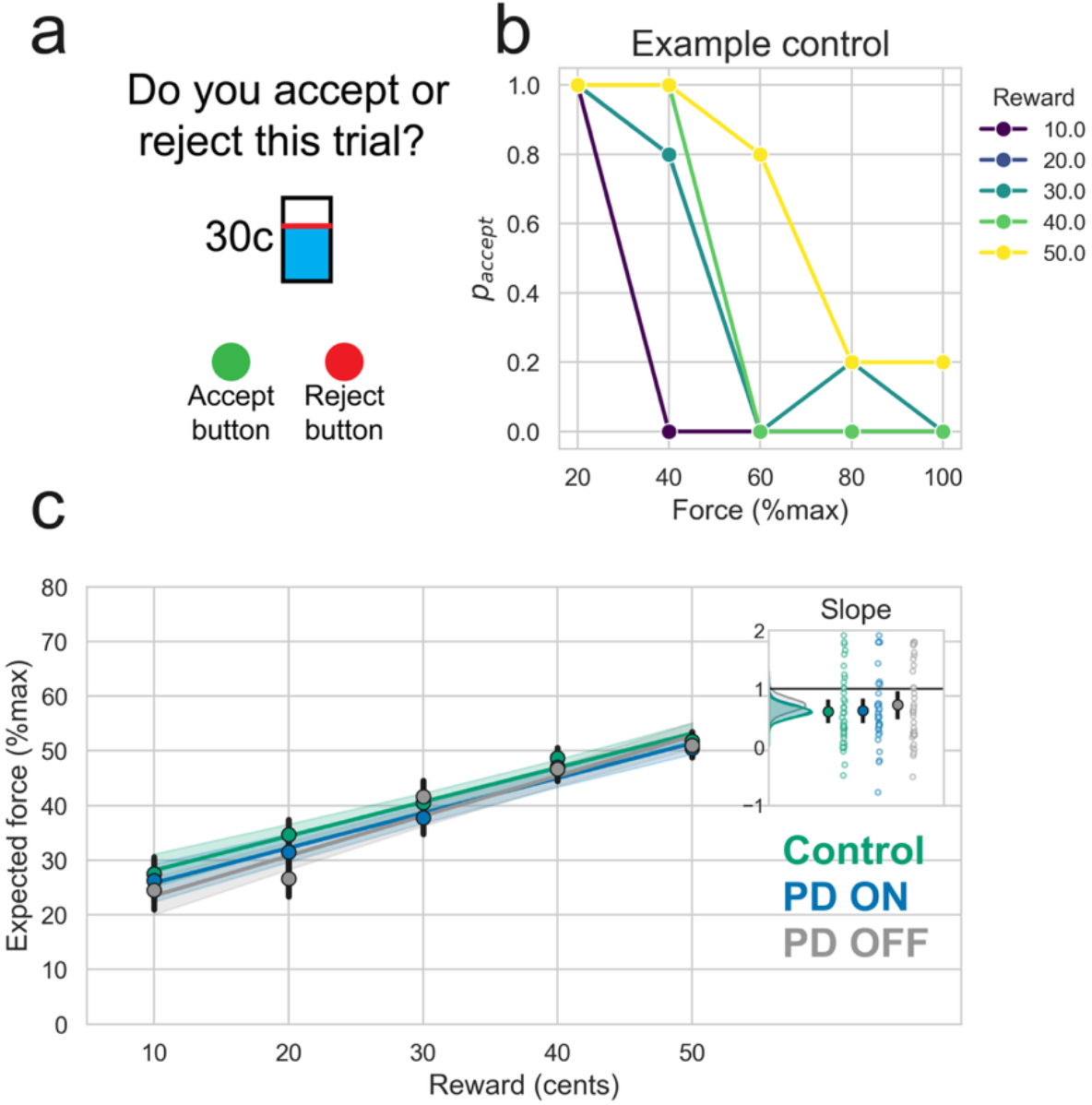
The mapping between effort and reward. **(a)** Participants decided if they were willing to produce a prospective isometric elbow extension force for a monetary reward by pressing either the accept or reject button. Each combination of 5 different force and 5 different reward levels were presented 5 times each. We calculated the expected force a participant was willing to perform at each reward level. Then, we regressed the expected force against reward for each participant. **(b)** Example control subject that was closest to median slope of all participants. Dots represent the probability of accepting the offer at each force and reward level. **(c)** Average empirical expected forces at each reward level (circles) plotted against the mean regressions. Error bars and shading represent 1 SEM. **(Inset)** The slope for the regression was our primary measure of the effort-reward mapping. Kernel density estimates of the posterior mean distributions are plotted alongside the empirical group means (filled circles) and individual datapoints (open circles). Error bars for these plots reflect the 95% HDIs for each posterior mean distribution

To quantify the mapping between effort and reward, we first calculated the expected force a participant was willing to accept at each reward level (equation 3, Methods). The forces an individual was willing to accept generally increased across reward levels (e.g., see Fig. 3b). To quantify this pattern, we regressed the expected forces onto the rewards for each participant. We estimated the effort-reward mapping as the slope of this regression since it reflects the change in the expected force a participant was willing to exert per unit change in reward. Steeper slopes indicated a greater willingness to exert more effort per unit change in reward. We found no differences in slopes between the PD ON and control groups (0.02 [−0.26 0.32], p_difference_ = 54.4%, d = 0.03), between the PD OFF and control groups (0.11 [−0.20 0.42], p_difference_ = 75.7%, d = 0.17) or between PD ON and PD OFF (−0.02 [−0.23 0.19], p_difference_ = 56.2%, d_z_ = −0.03). Hence, people with PD, regardless of medication status, exhibited a similar willingness to increase effort per unit change in reward as control participants. These results suggest that PD and dopaminergic medication do not bias the mapping between effort and reward.

### Cognitive perception of effort, but not reward sensitivity, predicts the willingness to produce effort in exchange for reward

In the analyses above, we directly examined how PD impacts each component of the effort-reward trade-off by separately assessing reward sensitivity, perceptions of effort, and the effort-reward mapping. However, a key question is how performance across these tasks is related. Thus, as a post hoc analysis, we examined whether reward sensitivity or the perception of effort influenced participants’ overall willingness to exert effort in exchange for reward (i.e., the acceptance rate during the effort-reward mapping task). We found that reward sensitivity (as measured by the PSE of the psychometric function fit to choice data in the reward sensitivity task) was not related to acceptance rates in the effort-reward mapping task for any group, as these effects were small and unreliable (PD ON z-scored beta = 0.07 [−0.34 0.44], p_zbeta>0_ = 63.2%; control zbeta = −0.05 [−0.42 0.29], p_zbeta>0_ = 62.1%; PD OFF zbeta = −0.05 [−0.33 0.45], p_zbeta>0_ = 59.8%).

For the perception of effort, lower force-matching intercepts (i.e., greater sensorimotor perceptions of effort) were associated with reduced acceptance rates during the effort-reward mapping task in the PD ON group (zbeta = 0.32 [−0.02 0.66], p_zbeta>0_ = 96.8%). However, the relationship was not reliable for controls (zbeta = 0.20 [−0.14 0.53], p_zbeta>0_ = 87.8%) or for PD OFF (zbeta = −0.06 [−0.41 0.29], p_zbeta>0_ = 63.3%). In contrast, slopes of the subjective effort ratings (i.e., greater cognitive perception of effort) were reliably related to lower acceptance rates during the effort-reward mapping task for both PD ON (zbeta = −0.48 [−0.80 −0.17], p_zbeta>0_ = 99.8%) and controls (zbeta = −0.31 [−0.62 0.01] p_zbeta>0_ = 97.2%). For the PD OFF group, this relationship was not reliable but remained in the same direction (zbeta = −0.19 [−0.55 0.17], p_zbeta>0_ = 85.2%). Together, these results suggest that perceptions of effort (particularly as reported through subjective ratings) influence how people decide to trade-off effort for reward.

## Discussion

The effort-reward trade-off is critical for motivating our actions, and impairments in this trade-off have been proposed to underlie low movement vigor (i.e., bradykinesia) in people with PD. We assessed three potential sources of impairment in the effort-reward trade-off in people with early-stage PD by individually interrogating reward sensitivity, perceptions of effort, and the mapping between effort and reward. We found that effort perception was disrupted in people with PD compared to neurotypical controls. In contrast, we found no difference between people with PD and controls in either reward sensitivity or the effort-reward mapping, nor was dopaminergic medication status associated with these factors. Overall, our results suggest that changes to the perception of effort, but surprisingly not reduced reward sensitivity, may be an important contributor to reduced vigor in people with PD.

Effort perception has largely been measured using one of two methods: having participants match a previously experienced force magnitude^38,41,47,60–63^ or having them rate the perceived effort exerted following force production^43–46^. While the former method tests sensorimotor-based perceptions^38–42^, the latter likely hinges on more cognitive processes. We observed that people with PD exhibited behavior consistent with perceptions of increased effort, relative to control participants, on both measures.

Disruptions to sensorimotor-based perceptions of effort could arise from a primary sensory impairment or impaired central processing of an ongoing motor command^38–42^. Several studies have reported reduced sensitivity to changes in limb position in people with PD, consistent with a primary somatosensory deficit (for review, see^48^). However, a broad somatosensory impairment implies that people with PD would have difficulty sensing changes in force magnitude, making it unlikely that they would reliably scale the relative magnitudes of their match forces with increasing initial force levels as we observed here. Moreover, we observed no impairment when people with PD produced initial forces with the more impaired arm but matched them with the less impaired arm (Supplemental Fig. 2), suggesting intact sensitivity to relative differences in the goal forces. Prior literature has also shown that people with PD can scale the relative magnitudes of isometric elbow flexion forces to accurately produce instructed percentages of their voluntary maximum force^64,65^. Note that, although we were constrained by the Kinarm robot’s maximum force output in our study, this prior work was not. This seeming discrepancy between the sensory and force production literature may stem from the fact that these sensations arise from distinct pathways: while the perception of limb position changes is thought to rely heavily on signaling from muscle spindles, isometric force perception is mediated by Golgi tendon organs (GTOs)^40^. PD has been shown to disrupt the fusimotor system and, in turn, spindle sensitivity^66,67^. It is currently unknown whether PD alters the processing of GTO signals, although the fact that people with PD can reliably scale their forces here and in prior studies suggests this is not the case.

Changes in sensorimotor effort perception could alternatively arise from disruptions in the central processing of the ongoing motor command. That PD participants were able to accurately produce the initial forces indicates that they can produce appropriate motor commands (i.e., there is no primary deficit in force production). However, people with PD produced match forces that were consistently lower than those of control participants, suggesting a systematic bias in the perceived effort associated with them. Prior studies of analogous force-matching paradigms have posited that effort perception may hinge on the correspondence between reafferent somatosensory feedback and predictive signals derived from motor commands. When asked to reproduce externally applied^42,68,69^ or visually calibrated^38,70^ forces using only reafferent somatosensation, neurologically healthy individuals exhibit systematic overshooting of the target force, which is thought to reflect compensation for a centrally mediated perceptual attenuation of the predicted component of the feedback signal. Notably, control participants showed this overproduction of matching forces in our study, while people with PD did not. PD has been shown to disrupt reafferent attenuation to a degree that increases with ratings of symptom severity and current dopaminergic medication dose^71,72^. Hence, disrupted effort perception in PD is consistent with an impairment in reafference processing.

When asked to retrospectively rate their perceived effort following match force production, people with PD, particularly when OFF their dopaminergic medication, reported greater exertion than control participants. This aligns with our interpretation that lower match forces reflected an increase in perceived effort and is consistent with prior work showing that people with PD rate the effort associated with everyday activities as greater than that of neurotypical controls^46^. We also found that effort perception ratings correlated with the overall willingness to exert effort in exchange for reward particularly for the PD ON and control groups, such that higher effort ratings were associated with reduced acceptance of effort-reward pairings in the trade-off task. While the trade-off between effort and reward also correlated with the sensorimotor perception of effort (i.e., match force magnitude) for the PD ON group, this relationship was unreliable in the control group. When considered together, these results suggest that the effort-reward trade-off may depend more on cognitively mediated than sensorimotor-based perceptions of effort.

While we found that the mapping between effort and reward was similar between people with PD and control participants, prior literature on the effort-reward trade-off in PD reports mixed results. Some studies found an altered mapping in people with PD compared to a control group^19,22,73^, but others did not^17,18^. A distinguishing feature of our study is that we did not require participants to immediately exert the forces they accepted after each decision, thereby reducing muscular fatigue throughout the task. Physical fatigue is known to increase the perception of effort^59^ and reduce the willingness to exert effort in exchange for reward^58^, and could account for the observed impairments in the effort-reward trade-off noted in prior studies. We also quantified the effort-reward mapping differently from prior studies, as the change in expected force a participant was willing to exert per unit change in reward rather than simply the overall willingness to exert effort for reward. These methodological differences could contribute to the mixed findings across studies.

We were surprised to find no evidence of reduced reward sensitivity in people with PD ON or OFF their dopaminergic medication relative to controls. We assessed reward sensitivity using a utility task commonly employed in the behavioral economics literature, which quantifies how much a given monetary reward is worth to an individual (i.e., its subjective rather than objective value)^32,33,53,54^. Our findings conflict with prior studies that have observed changes in reward sensitivity in people with PD relative to control participants using the same utility task^34–37^, and with physiological studies that have linked dopamine neuron activity to subjective reward value^74–77^. However, prior studies of reward sensitivity in people with PD typically provided reward feedback immediately after each decision, which has been shown to bias decisions in utility tasks and violates the assumption that these tasks probe a stable measure of reward sensitivity^78,79^. To avoid this confound, we did not provide decision feedback in our study. Notably, one prior study that withheld decision feedback in a similar manner also found that people with PD exhibited reward sensitivity similar to that of control participants^21^.

How can we reconcile our results with the well-established connection between dopamine signaling and reward during reinforcement learning^80–83^, as well as findings of dopamine-dependent impairments in reinforcement learning observed in people with PD^29–31,84–86?^ First, we note that our reward sensitivity task did not involve learning: each decision was independent, and no decision feedback was available as a teaching signal to update the value assigned to subsequently presented certain and gamble options. Instead, our reward-sensitivity task focused on evaluating the perceived current value of a familiar reward. Dopamine signaling has been predominantly linked to reward in the context of learning, with some recent work even suggesting that dopamine can serve as a value-free teaching signal^87^. In contrast, dopamine signaling might not be as critical for recalling the value assigned to already learned rewards^88–90^. In this way, findings of dopamine-dependent reinforcement learning impairments in people with PD may reflect a problem with the learning process rather than baseline reward sensitivity. Additionally, we note that a significant proportion of studies relating dopamine and reward focus on the ventral tegmental area and its connections to the prefrontal cortex^75–77,80–82^. Conversely, PD tends to affect dopaminergic neurons in the substantia nigra and their connections to regions of the basal ganglia that communicate with motor cortex, especially early in disease progression^24,91–97^. Hence, it is an oversimplification to conclude that, because dopamine signaling has been linked to reward and dopamine-producing neurons degenerate in PD, people with PD must have an impairment in reward sensitivity. Critically, people in earlier stages of PD still exhibit a decrease in movement vigor that is thought to reflect an underlying imbalance in the effort-reward trade-off. Our findings suggest that, for these individuals, bradykinesia is driven by increased effort perception rather than decreased reward sensitivity.

Like prior studies, the current study relied on isometric force production as a proxy for effort and monetary reward as a proxy for motivation. However, this may not fully capture real-world motivation or effort in PD. Similarly, our assessment of reward sensitivity addressed confounds in prior literature by isolating the parameter from learning, but we note that reward valuation and learning from reward often co-occur in everyday contexts, and this may also limit the ecological validity of our results. Finally, our sample of people with PD was mostly in the early stages of the disease process as measured by the Hoehn and Yahr scale. While we did not observe relationships between motor symptom severity (UDPRS motor score) and any of the primary outcome measures for the sample of participants in the current study, a larger and more inclusive sample of individuals with PD that could be stratified by disease stage would better enable us to examine the effect of dopamine medication across different stages of the disease process.

In summary, we observed that people with early-stage PD show a selective increase in the perceived effort associated with motor output relative to neurotypical controls, without a concomitant change in reward sensitivity or the mapping between effort and reward. We posit that biased effort perception underlies previous findings of altered effort-reward trade-offs in people with PD and contributes to the reduction in movement vigor (i.e., bradykinesia) commonly observed in this population. Indeed, people with PD frequently produce slower movements than neurotypical individuals unless sufficiently motivated^14,15^, and even then, their movement vigor does not respond to reward as effectively as in neurotypical controls^3^. Our results imply that reduced vigor and impaired scaling of vigor with reward in PD stem from an underlying disruption of the perception of effort associated with more vigorous movements. Notably, this aligns with conventional rehabilitation approaches for PD, which aim to recalibrate people’s perception of their own movements rather than providing greater incentives (e.g.,^98^). Our results suggest that further exploration of this approach may be beneficial when designing future treatments for people with PD.

## Methods

### Participants

We recruited 44 participants with a diagnosis of idiopathic mild to moderate PD (Hoehn and Yahr^99^ <=3) and 40 neurotypical controls matched for age (± 3 years) and sex. Participants in the PD group were excluded if they had a deep brain stimulator implant, any neurologic diagnoses aside from PD, or any psychiatric diagnoses unrelated to PD. Controls were excluded if they had a diagnosis of any neurologic or psychological disorders. Participants were excluded if they had any significant orthopedic problems that could interfere with testing or significant cognitive impairment, defined as a score of <20 on the Montreal Cognitive Assessment (MOCA^100,101^). Seven participants in the PD group were excluded due to an inability to complete all tasks (n=3), impaired cognition as measured by the MOCA (n=2), or were subsequently diagnosed with a Parkinson’s plus syndrome after testing (n=2). One participant in the control group was removed because they were unable to complete the tasks. This left a final sample of 37 participants with PD (11 female) and 39 controls (13 female). There were no differences in age between the final samples of PD and control participants (mean difference [95% HDI] = 0.08 [−3.2 3.5], p_difference_ = 51.7%, d = 0.01). Participant demographics are summarized in Table 1. All participants were paid $20/hour for study participation with the opportunity to earn additional bonus payments based on the rewards earned during the experiment. All participants provided written informed consent prior to the study, and all experimental procedures were approved by the Thomas Jefferson University Institutional Review Board (IRB-2020-407)

**Table 1:**
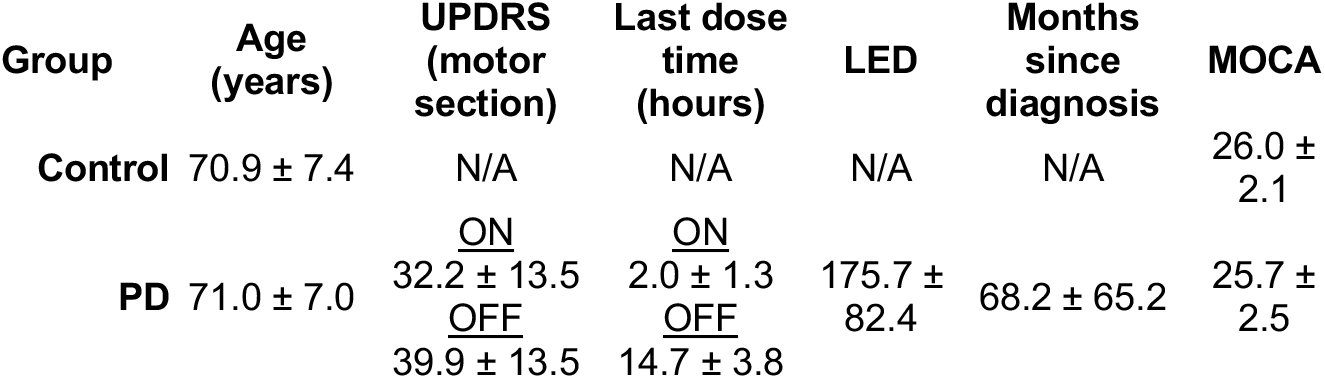
Demographics. Mean ± 1 SD. UPDRS: Unified Parkinson’s disease rating scale, LED: Levodopa equivalent dose, MOCA: Montreal Cognitive Assessment. A detailed breakdown of the PD group demographics is included in the Supplementary Table 1.

### Study procedure

Using a Kinarm Exoskeleton Robot (Kinarm, Canada), participants completed three tasks in the following order over a single experimental session: the force-matching, reward sensitivity, and effort-reward mapping tasks (detailed below). This fixed task order was chosen to mitigate muscular fatigue. All tasks were custom-programmed in Simulink (version R2015a, Mathworks, United States) and analyzed using MATLAB (v2022a) for preliminary analysis. Simulink code to run the tasks on the Kinarm is available on the lab GitHub page (force-matching task: https://github.com/CML-lab/KINARM_Iso_Force_Matching; reward sensitivity task: https://github.com/CML-lab/KINARM_Utility_Test; effort-reward mapping task: https://github.com/CML-lab/KINARM_Effort_Reward_Mapping). The remainder of the analyses were performed in Python (v3.13.5). Each task involved either isometric force production or single-arm reaching. Because motor symptoms are typically lateralized in PD, participants with PD performed each task with their self-reported, more-affected arm. For the PD group, the same arm was used for the ON and OFF sessions. Each control participant performed the tasks using the arm that matched the dominance or non-dominance of the tested arm of the PD participant to whom they were matched. In terms of dopamine-affecting medications, 34 participants with PD were taking a carbidopa/levodopa medication, of whom 5 were also taking a dopamine agonist. Of the remaining 3, one participant was only taking MOA-B and COMT inhibitors, one was only taking a dopamine agonist, and the other was not taking any dopamine medication. To determine the impact of medication status on the individual components of the effort-reward trade-off, 32 of the participants with PD agreed to be tested in both the ON and OFF medicated state in separate sessions. We counterbalanced the order of the ON versus OFF sessions as much as possible (18/32 completed the ON test first). Three participants with PD completed only the OFF test: one was not on any medication, and the other two were unable to return for the ON test. For those who completed both the ON and OFF sessions, the second session was performed at least 18 days after the first session (mean = 50.1, SD = 24.8 days). For the ON test, participants were tested an average of 2.0 (SD = 1.3) hours after their most recent dose of dopaminergic medication. For the OFF test, participants skipped a single morning dose of dopaminergic medication. They were tested an average of 14.7 (SD = 3.8) hours after their last dose. Motor symptom severity as measured by Part III of the Movement Disorders Society Unified Parkinson’s Disease Rating (UPDRS)^102^ was lower for people with PD during the ON condition compared to OFF (−8.3 [−11.4 −5.3], p_difference_ = 100.0%, d = −1.02). No reliable relationship was observed between UPDRS scores and the primary outcome measures for any of our tasks.

### Reward sensitivity task

#### Procedure

To assess reward sensitivity, we adapted a utility task from the prior literature^32,33,53,54^ (Fig. 1a). Participants were given the choice between a guaranteed monetary reward and a 50/50 chance of winning a different monetary reward (i.e., the certain and gamble options, respectively). The two options were displayed as square targets (8 cm width) on the Kinarm display screen. Direct vision of the arm and hand was occluded throughout the task. Participants were provided visual feedback of their hand movement in the form of a cursor (white circle with 0.5 cm radius) aligned to the position of the index fingertip on the Kinarm display screen.

A trial began with the participants moving the cursor into a green start circle (2 cm radius). After a random delay (uniformly sampled between 500 and 700ms), two square targets appeared on the screen with the certain and gamble options displayed inside. After another 1800ms delay, an auditory “go” signal sounded, cueing participants to indicate their preferred option by reaching as quickly and accurately as possible to the corresponding square target. If a reach was initiated before the go signal, the square targets were extinguished, and the trial was removed from analysis (4.5% of trials). Keeping these trials in the analysis by inferring the choice based on hand position proximity to the target either at the beginning or at the end of the trial did not change the results. The trial ended once the cursor remained inside a square target for 500ms, or if the cursor did not successfully intersect a target within 6 seconds after the go signal sounded (less than 0.1% of trials).

Participants completed 5 blocks of 27 combinations of certain (10-50 cents) and gamble (10-100 cents) reward magnitudes for a total of 135 trials. The certain and gamble pairs were randomly presented during each block. For half these trials, the certain target would appear on the left side of the screen, and for the other half, it would appear on the right side of the screen. To reduce fatigue, a rest break was provided after the second and fourth blocks. Neither the result of the gamble nor the running total of their earnings was displayed to the participants, as this may have encouraged people to change their behavior in light of gamble outcomes observed on prior trials. Participants were also not informed of the number of trials remaining in the task. Prior to the task, participants were informed of the 50/50 probability of the gamble options, and that a portion of their earnings would be added to their study payment. To ensure they understood the task and the instructions, participants completed 5 practice trials before performing the task.

#### Data analysis

From the data collected for this task, we first calculated the expected values (EV) of the certain and gamble options on each trial:

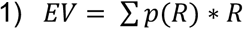

where R represents the reward magnitude and p represents the probability. We then subtracted the expected value of the certain option from the expected value of the gamble option, and recorded the number of times the gamble option was selected at each EV difference level. We fit a logistic function with two free parameters to each participant’s choice data using maximum likelihood estimation:

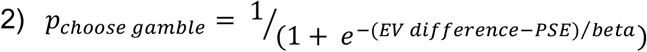

The point of subjective equality (PSE) represented the inflection point in the curve or the difference in expected value where the participant was equally likely to select the certain and gamble options. The beta parameter reflects noise in the decisions. To ensure stable fits for the parameters, we used 60 initializations sampled from normal distributions centered on empirical estimates of the PSE and beta values. We chose this option because fitting a more traditional utility function to the data^32,33^ resulted in less stable fits, but had no impact on the overall results. PSE was used as the primary outcome variable as it reflected participants’ general reward sensitivity. More positive PSE values indicated that participants required the gamble option to have greater expected values to select it. Conversely, more negative PSE values indicated that participants did not require the gamble option to have greater expected values to overcome the uncertainty associated with choosing it. The fitted logistic function was also used to quantify the just noticeable difference (JND), defined as the change in expected value difference between the 75^th^ and 25^th^ percentiles. The JND reflected the minimum required change in reward value for a participant to reliably alter their choice behavior. Greater JNDs indicated a lower sensitivity to changes in the EV difference between the certain and gamble options.

Participant data was removed from this analysis if their choice data could not be reliably fit with a logistic function, i.e., if their probability of choosing the gamble option was never above p=0.75 or below p=0.25 across all levels of expected-value difference. According to this threshold, we removed 4 control participants, 2 PD participants for both ON and OFF conditions, 3 PD participants for the ON condition only, and 3 PD participants for the OFF condition only. To confirm that removing these individuals did not bias our results, we also calculated the probability of choosing the gamble option on trials where the expected values of the two options were equal as a secondary measure of reward sensitivity.

Asking participants to move as quickly and accurately as possible after the go signal allowed us to also assess the vigor of their reaches during this task. Specifically, we calculated the peak velocity of the reach to the target. Because peak velocity is modulated by movement amplitude, and the large target allowed for a range of reach amplitudes, we normalized peak velocity by the amplitude of each reach. To determine if reward magnitude impacted the vigor of these reaching movements^3,103^, we estimated the slope of a linear regression for each individual, with the total reward on offer in each trial as the predictor and normalized reach velocity as the outcome (see Supplemental Results).

### Force matching task

#### Procedure

To assess sensorimotor and cognitive perceptions of effort, participants performed a force-matching task. Direct vision of the arm and hand was occluded throughout the task. This task involved an isometric elbow extension force, chosen because people with PD have greater difficulty performing extension movements than flexion movements^57^. First, we assessed participants’ maximum voluntary contraction (MVC) of elbow extension by asking them to extend their elbow and flex their triceps as hard as possible, keeping their shoulder and wrist relaxed. The average of 3 measurements was taken as the MVC. However, we were limited by the ability of the Kinarm robot’s motors to produce forces of equal magnitude to those produced by participants to keep the arm from moving. To stay within the limits of the motors, the max force was set to 20 N for anyone whose actual MVC exceeded 20 N (max force = 20 N in 27 PD ON, 27 PD OFF, and 38 control participants).

For each trial of the force-matching task, participants first produced an initial force with the aid of visual feedback, which consisted of a rectangular bar on the Kinarm display (Fig. 2a). On each trial, the target force level was mapped to the top of the bar, while the visual height of the bar remained constant across trials. The fill of the bar increased in proportion to the magnitude of force produced. Participants were instructed to “fill the bar” without exceeding the bar’s height. The initial force was considered successful if participants met one of two conditions: they maintained an isometric force magnitude within ±1 N of the target force for 1 s, or 5 s passed after initiating a minimum required force level (defined as the target magnitude minus 5% of the maximum force). Following successful completion of the initial force, participants were instructed to rest for 2 s before producing the match force – i.e., reproducing and maintaining the perceived initial force magnitude without visual feedback until the trial ended. Participants were provided a maximum of 5 seconds to produce the match force. Following production of the match force, participants were instructed to rate how effortful they perceived it to be on a scale of 0 to 10, with 10 representing the greatest possible effort and 0 representing no effort. Participants were tested on four different goal forces, 20%, 40%, 60%, and 80% of their max force, presented randomly within four blocks for a total of 16 trials. One PD participant performed an earlier version of this task in which they were presented with an additional target force of 50% of max, with all five force levels repeated three times each for a total of 15 trials.

Participants completed two runs of the force-matching task. In the first run, participants performed the initial and match forces using the same arm. In the second run, match forces were produced with the arm opposite the one used to the produce the initial forces (see Supplemental Results). Hence, for the PD groups, the first run tested effort perception in the more affected arm (data presented in the Results), and the second run tested effort perception as reported by the less affected arm (data presented in the Supplemental Results). Aside from the change in arm during the match force, the procedure remained the same for both runs of the task.

To reduce fatigue, a brief rest break was provided halfway through each run of the force matching task. To ensure that participants understood the procedure, they performed three practice trials prior to the start of the task, during which they received visual feedback during both the initial and match force production phases. The target forces practiced here were 25%, 50%, and 75% of the max force, presented in a fixed order.

#### Data analysis

We calculated the initial and match forces as the mean elbow extension force produced when the force level reached a plateau (i.e., the longest stable period of force production). These plateaus were assigned manually in Matlab by visually inspecting the force trace for each trial, as movement impairments in some of the participants with PD (e.g., due to tremor) prevented the use of a purely automated method. For all analyses, we removed trials in which the initial force never reached the goal threshold, defined as the goal force minus 5% of the max force (0.8% of all trials). Retaining these trials in the analysis did not change the results.

Data analysis comprised simple linear regressions. Goal forces, initial forces, match forces, and subjective report data were all mean-centered, allowing us to interpret regression intercepts as the outcome relative to the mean force level. We quantified initial force performance by regressing the initial forces against the goal forces for each participant. We quantified match force performance by regressing the match forces against the initial forces for each participant. Subjective effort ratings were quantified by regressing the subjective ratings of match forces against the initial forces for each participant. In all cases, a slope and intercept term from each regression was extracted for statistical analysis.

### Effort-reward mapping task

#### Procedure

To assess the mapping between effort and reward, participants performed a decision-making task adapted from the prior literature^17,18^. The task had participants indicate their willingness to produce varying magnitudes of force in exchange for varying monetary rewards. The forces corresponded to isometric elbow extension forces, presented as a rectangular bar on the Kinarm display screen with a red line inside it (Fig. 3a). The bar was calibrated so that the top corresponded to 100% of the max force determined in the force-matching task, with the red line indicating the offered force magnitude in each trial. Choice options varied between five force levels (20%, 40%, 60%, 80%, and 100% of the max force) and five reward levels (10¢, 20¢, 30¢, 40¢, and 50¢). Each combination of force and reward was presented randomly 5 times, resulting in a total of 125 trials.

Participants used buttons fixed to the left-hand trough of the Kinarm to accept or reject each presented offer. To mitigate the chances of muscular fatigue impacting choice behavior^58,59^, participants were not required to produce the offered force immediately upon accepting. Instead, participants were instructed that the experimenter would randomly select five trials from the pool of accepted offers whose force levels would need to be produced at the end of the task. Participants were informed that if they accepted fewer than five offers, the experimenter would randomly select from the pool of all possible offers. Participants were informed that successful completion of the five selected force levels would result in the corresponding rewards being added to their study payment. Prior to the task, participants were informed of the minimum and maximum reward and force levels.

To ensure familiarization with each force level offered, participants completed a block of trials requiring them to produce each force level (except for the 100% level) before beginning the decision-making task. No rewards were offered during this familiarization block. Additionally, participants completed 2 rounds of practice with the decision trials to familiarize them with the prospective nature of the task.

#### Data analysis

Using participants’ choice behavior, we calculated the probability of acceptance for each combination of effort and reward values. Using these probabilities, we calculated the expected force (EF) that people were likely to accept at each reward level:

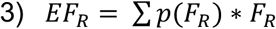

Where F = force magnitude, R = reward level, and p = probability. We next performed a linear regression for each participant with reward as the predictor and expected force as the outcome variable. We used the slope of this regression to quantify the mapping between effort and reward, since it reflects the change in expected force that someone is willing to accept for a unit change in reward. Higher slopes indicate that an individual was more willing to exert additional effort for a unit change in reward, while lower slopes indicate they were less willing to exert additional effort for a unit change in reward.

#### Statistical analysis

All statistical analyses were performed in Python using the PyMC^104^ (v5.23.0) and Bayes Toolbox^105^. Analysis code used for generating figures and conducting statistical analyses is available on https://osf.io/krw49.

We performed separate Bayesian t-tests for the between-subject (PD ON vs controls and PD OFF vs controls) and within-subject (PD ON vs. PD OFF) comparisons for each outcome of interest^52^. The Bayesian t-tests for the between-subjects comparisons assumed the data (e.g., intercept, slope, or PSE) from an individual participant was generated from one of two t-distributions, depending on the group (PD ON or control, and PD OFF or control). The distributions for each group had a mu and a sigma parameter, and a single nu parameter was shared between the groups. For the within-subjects Bayesian t-test, we assumed the data were generated from a single t-distribution of within-subject differences in the outcome of interest. This distribution had a single mu, sigma, and nu parameter. For all comparisons, we were most interested in the posterior distribution of the mu parameters. We set a wide and uninformative prior on each of the model parameters to avoid undue influence on the posterior^51^. We used Bayes’ rule to estimate the posterior distribution of these parameters by combining the data with the prior. To estimate the joint posterior distribution, we used Markov Chain Monte Carlo sampling with 4 chains, 10,000 samples, and 2,000 tuning steps. We visually inspected the trace and the posterior predictive checks of the model against the data to ensure a reasonable model fit^51^.

For the between-subject comparisons, we subtracted the posterior distributions of the mu parameters. This distribution reflected the difference in means that most likely generated the data. The same interpretation can be applied to the posterior distribution of differences for the within-subject comparisons. For each of these posterior distributions, we report the mean and 95% high-density interval (HDI). The HDI is defined as the narrowest span of credible values that contains 95% of the distribution (i.e., the true difference falls within this range with 95% certainty). We use this range of values to infer the magnitude of the effect. We also calculated the percentage of the posterior distribution of differences that fall on one side of 0 (p_difference_).

This percentage reflects the probability (i.e., reliability) of a difference. We always interpret this probability as higher values indicating a greater probability of a true difference. We considered both the magnitude and reliability of the effect together when interpreting our results, with p_difference_ values >95% being considered sufficiently reliable. Lastly, we report an estimate of standardized effect size, calculated from the posterior distribution of mean differences and standard deviations. This calculation is equivalent to Cohen’s d and d_z_ for the between and within subjects comparisons, respectively. We report the mean of the posterior distribution of the standardized effect sizes.

To make inferences about the similarity between groups specifically for the reward-sensitivity analysis, we calculated a Bayes factor ^55,56^. This approach has greater power than alternative equivalence tests (e.g., a Region of Practical Equivalence) for the sample size in the current study^106^.

To make inferences regarding the relationship between outcomes, we employed Bayesian regressions. Here, we assumed that each outcome (y) for each participant (i) was generated from a t-distribution with a mu parameter, a sigma parameter, and a nu parameter. In this model, the mu parameter depends on the size of a predictor variable (x) and two beta parameters (β_0_ and β_1_) according to the following equations:

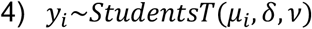

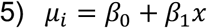

For this analysis, we report the standardized beta coefficients by performing the analysis on z-scored data (zbeta). We calculated the posterior in the same manner as the Bayesian t-tests, along with the 95% HDI and the percent of the posterior distribution of betas that are on one side of 0 (p_zbeta>0_), to determine the magnitude and reliability of the effect, respectively.

## Supporting information

Supplemental Results

## Acknowledgements

This project was supported by the Klein Family Parkinson’s Rehabilitation Center (JMW and ALW), a MossRehab Peer Review Committee grant (PRC FY20-1) to ALW, and the National Science Foundation (BCS2444305) to AST. We would like to thank Sky Yallof and Luke Carter for their help with data collections.

