## Supplemental Results for "Impaired effort perception, not reward sensitivity, impacts the effort-reward trade-off in Parkinson’s disease"

#### Demographics table for participants with Parkinson's disease

| Participant ID | Age (years) | Sex | Months since diagnosis | Handedness | More affected arm* | MOCA | H&Y | UPDRS Section III | Hours since last medication dose | LED |
|---|---|---|---|---|---|---|---|---|---|---|
| **PD-001** | 66 | F | 135.2 | Right | Right | 28 | 2 | ON = 21<br>OFF = 25 | ON = 1<br>OFF = 15 | 133 |
| **PD-002** | 70 | M | 136.2 | Right | Left | 29 | 2 | ON = 10<br>OFF = 23 | ON = 0.5<br>OFF = 17.5 | 343 |
| **PD-003** | 67 | M | 74.0 | Right | Right | 26 | 2 | ON = 15<br>OFF = 16 | ON = 1.5<br>OFF = 17.5 | 150 |
| **PD-004** | 74 | M | 72.0 | Right | Right | 29 | 2 | ON = 40<br>OFF = 36 | ON = 0.5<br>OFF = 17.5 | 125 |
| **PD-005** | 75 | M | 94.4 | Right | Right | 24 | 2 | ON = 35<br>OFF = 38 | ON = 1.5<br>OFF = 17.5 | 200 |
| **PD-006** | 72 | F | 267.0 | Right | Left | 24 | 3 | ON = 40 | ON = 1.5 | 160 |
| **PD-007** | 80 | M | 32.4 | Right | Left | 25 | 2 | ON = 25<br>OFF = 28 | ON = 6<br>OFF = 22 | 100 |
| **PD-008** | 73 | M | 35.2 | Right | Left | 28 | 2 | ON = 34 | ON = 0.5 | 250 |
| **PD-009** | 71 | F | 44.6 | Right | Right | 26 | 2 | ON = 35<br>OFF = 43 | ON = 4<br>OFF = 15.7 | 100 |
| **PD-010** | 75 | M | 77.2 | Right | Right | 24 | 2 | ON = 43<br>OFF = 46 | ON = 2<br>OFF = 12.2 | 200 |
| **PD-011** | 69 | M | 76.0 | Right | Left | 30 | 2 | ON = 23<br>OFF = 32 | ON = 2.3<br>OFF = 16.2 | 300 |
| **PD-012** | 59 | M | 40.8 | Right | Right | 28 | 2 | ON = 26<br>OFF = 35 | ON = 1.9<br>OFF = 22.1 | 75 |
| **PD-013** | 65 | M | 81.5 | Left | Left | 21 | 2 | ON = 42<br>OFF = 55 | ON = 3.8<br>OFF = 25.2 | 100 |
| **PD-014** | 78 | F | 48.2 | Right | Left | 26 | 2 | ON = 24<br>OFF = 43 | ON = 3.1<br>OFF = 12.2 | 175 |
| **PD-015** | 79 | M | 67.1 | Right | Right | 26 | 2 | ON = 34<br>OFF = 42 | ON = 1.3<br>OFF = 14.9 | 100 |
| **PD-016** | 71 | M | 8.7 | Right | Left | 20 | 2 | ON = 35<br>OFF = 39 | ON = 1.6<br>OFF = 6 weeks† | 200 |
| **PD-017** | 69 | F | 11.4 | Left | Right | 25 | 2 | ON = 16<br>OFF = 30 | ON = 1.5<br>OFF = 16.2 | 100 |
| **PD-018** | 82 | M | 48.1 | Right | Right | 22 | 2 | OFF = 21 | OFF = 12 | 75 |
| **PD-019** | 82 | M | 50.5 | Right | Left | 27 | 2 | ON = 36<br>OFF = 38 | ON = 1.8<br>OFF = 14.0 | 350 |
| **PD-020** | 67 | F | 68.1 | Right | Left | 24 | 2 | ON = 29<br>OFF = 49 | ON = 2<br>OFF = 13.4 | 133 |
| **PD-021** | 67 | M | 20.5 | Right | Right | 27 | 2 | ON = 35<br>OFF = 35 | ON = 2.5<br>OFF = 11 | 275 |
| **PD-022** | 69 | M | 36.9 | Right | Right | 26 | 2 | ON = 25<br>OFF = 28 | ON = 0.85<br>OFF = 12.5 | 275 |
| **PD-023** | 68 | M | 98.1 | Right | Left | 27 | 2 | ON = 28<br>OFF = 39 | ON = 1<br>OFF = 10.0 | 167 |
| **PD-024** | 73 | M | 349.9 | Right | Right | 25 | 2 | ON = 54<br>OFF = 63 | ON = 1.3<br>OFF = 12 | 207 |

| | | | | | | | | | | |
|---|---|---|---|---|---|---|---|---|---|---|
| **PD-025** | 69 | M | 25.4 | Right | Right | 27 | 2 | ON = 23<br>OFF = 44 | ON = 2.0<br>OFF = 12 | 195 |
| **PD-026** | 87 | F | 94.6 | Right | Right | 23 | 2 | ON = 56<br>OFF = 44 | ON = 3<br>OFF = 5.5 | 175 |
| **PD-027** | 63 | M | 32 | Right | Left | 27 | 2 | ON = 48<br>OFF = 66 | ON = 1.8<br>OFF = 11.7 | 100 |
| **PD-028** | 61 | F | 12.2 | Right | Right | 23 | 3 | ON = 67<br>OFF = 69 | ON = 1.1<br>OFF = 15.5 | 100 |
| **PD-029** | 63 | F | 19.5 | Right | Right | 28 | 2 | ON = 18<br>OFF = 16 | ON = 2<br>OFF = 11.4 | 220 |
| **PD-030** | 69 | M | 24.4 | Right | Left | 27 | 2 | ON = 33<br>OFF = 50 | ON = 6<br>OFF = 16.2 | 100 |
| **PD-031** | 77 | M | 26.8 | Right | Left | 29 | 1 | ON = 23<br>OFF = 50 | ON = 1.8<br>OFF = 16.5 | 200 |
| **PD-032** | 74 | F | 112.1 | Right | Right | 28 | 0 | ON = 10<br>OFF = 27 | ON = 1.1<br>OFF = 11.5 | 370 |
| **PD-033** | 74 | M | 36.2 | Right | Right | 21 | 2 | ON = 59<br>OFF = 68 | ON = 3.7<br>OFF = 14.6 | 150 |
| **PD-034** | 60 | M | 16.1 | Right | Left | 27 | 2 | ON = 24<br>OFF = 42 | ON = 1.4<br>OFF = 11.8 | 100 |
| **PD-035** | 66 | F | 82.3 | Right | Right | 28 | 2 | ON = 30<br>OFF = 33 | ON = 1.1<br>OFF = 17 | 200 |
| **PD-036** | 60 | M | 35.3 | Right | Right | 23 | 2 | OFF = 42 | OFF = n/a‡ | 0 |
| **PD-037** | 84 | F | 106.9 | Right | Right | 23 | 3 | OFF = 46 | OFF = 15.8 | 200 |

**Supplementary table 1.** List of demographic variables for each participant with PD. MOCA = Montreal cognitive assessment; H&Y = Hoen and Yahr score, taken during their first test date; UDPRS = Unified Parkinsons Disease Ratings Scale Motor Scale; LED = Levodopa Equivalent Dose

* Self-reported

† This individual was taking their prescribed medication (Amantadine) for the ON test but stopped taking it for about 6 weeks before the OFF test - this value was not included in the summary statistics.

‡ This individual was not on any medication

### Movement vigor during the reward sensitivity task

We assessed the vigor of the reaching movements during the reward sensitivity task. In this task, participants were instructed to reach as quickly and accurately as possible to the target of their choice immediately following a go signal, which sounded 1800 ms after the targets appeared. Since participants could stop anywhere within the rather large targets (8cm squares), we normalized peak velocity by movement amplitude as our measure of vigor. For each participant, we averaged the normalized peak velocity across all valid trials. We found that people with PD moved slower compared to controls (PD ON vs controls mean difference [95% HDI] = -0.42 [-0.71 -0.12], $p_{difference}$ = 99.7%, d = -0.69). Additionally, people with PD moved slower when OFF their dopamine medication compared to ON (-0.18 [-0.28 -0.07], $p_{difference}$ = 99.8%, $d_z$ = -0.70), and PD OFF moved slower compared to controls (-0.59 [-0.88 -0.29], $p_{difference}$ = 100.0%, d = -0.96). This confirms that within our sample, people with PD indeed moved with reduced vigor compared to controls, and a lack of dopamine exacerbated this effect.

Prior research has demonstrated that movement vigor increases alongside the value of the offered reward[1] and potentially that movement vigor reflects subjective value of monetary reward[2,3]. Therefore, we also performed an analysis to determine how reward magnitude impacted movement vigor on each trial (Fig. S1). For each participant, we regressed the z-scored, normalized vigor against the z-scored total reward magnitude offered on each trial. The slope of this regression reflects the impact of reward on movement vigor. If people with PD were less sensitive to differences in reward value, they should exhibit smaller slopes for this regression. We found that the slopes were positive across the three groups (control mean [95% HDI] z-scored beta = 0.03 [0.02 0.05], $p_{zbeta>0}$ = 100.0%; PD ON = 0.03 [0.01 0.05], $p_{zbeta>0}$ = 99.9%; PD OFF = 0.03 [0.01 0.04], $p_{zbeta>0}$ = 100.0%), thus replicating the prior work that movement vigor increases with greater reward magnitude. However, we observed no differences between the magnitudes of these slopes for any comparison (PD ON vs control = -0.001 [-0.02 0.02], $p_{difference}$ = 51.9%, d = -0.01; PD ON vs PD OFF = 0.002 [-0.02 0.02], $p_{difference}$ = 56.7%, $d_z$ = 0.03; PD OFF vs control = -0.01 [-0.03 0.01], $p_{difference}$ = 80.4%, d = -0.21). Therefore, while people with PD did reach with reduced vigor – an effect that was exacerbated when OFF dopamine medication – their vigor was not modulated differently by monetary reward. This provides additional evidence against the hypothesis that reward sensitivity is impaired in people with PD.

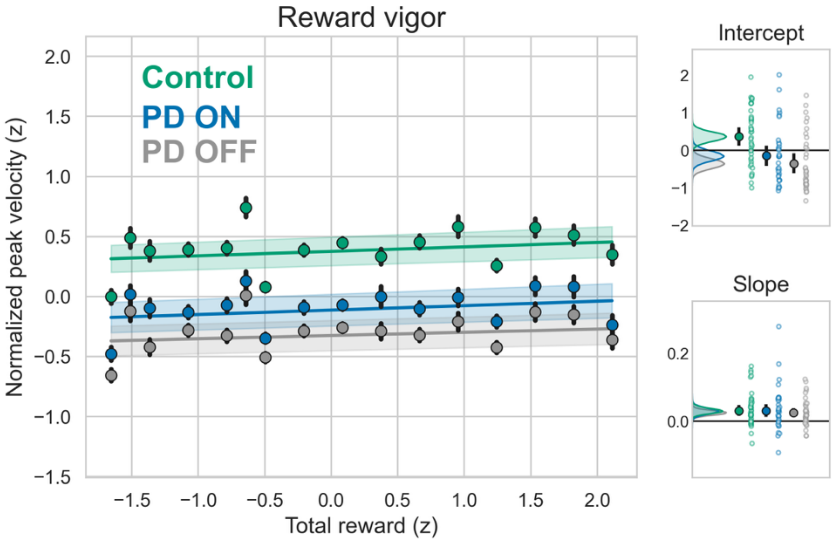


**Figure S1. Reward Vigor.** The main plot depicts the group averaged normalized peak velocity plotted at each reward level (both z-scored). Each dot represents the mean of each group, and error bars are 1 standard error of the mean. Lines and shading represent the mean regression line and 1 standard error of the mean, respectively. Insets show the intercept and slopes of the participant level regressions. Kernel density estimates of the posterior mean distributions plotted alongside the empirical group means (filled circles) and individual datapoints (open circles). Error bars for these plots reflect the 95% HDIs for each posterior mean distribution.

### Force matching task for the less affected arm

Participants completed two runs of the force-matching task (except for 1 individual in the PD group who was unable to complete the opposite arm task due to a recent shoulder surgery in the less-affected arm). Both runs were the same aside for the arm used to produce the match force. In the first run, participants performed the initial and match forces using the same arm. In the second run, match forces were produced with the arm opposite the one used to the produce the initial forces. Since the severity of motor symptoms in PD tends to be lateralized, the first run tested effort perception in the more affected arm while the second run tested effort perception in the less affected arm. Here we report results from the opposite-arm run, which were analyzed in the same manner as the same-arm results as described in the Methods.

We first assessed performance on the initial force (Fig. S2a). We regressed each participant's mean-centered initial forces against the respective mean-centered goal forces. The resulting intercepts reflected the initial force (relative to the mean) produced at the average goal force level, and the slope reflected the change in the initial force across the goal force levels (Fig. S2c). There were no differences in either the intercepts (PD ON vs controls = -0.2% max force [-0.6 0.2], $p_{difference}$ = 87.5%, d = -0.30; PD ON vs PD OFF = 0.2% max force [-0.1 0.5], $p_{difference}$ = 88.5%, $d_z$ = 0.25) or slopes (PD ON vs controls = -0.001 [-0.01 0.01], $p_{difference}$ = 55.0%, d = -0.03; PD ON vs PD OFF = 0.01 [-0.01 0.02], $p_{difference}$ = 87.7%, $d_z$ = 0.23). While the PD OFF condition demonstrated lower intercepts during the initial forces compared to controls (-0.4% max force [-0.8 -0.04], $p_{difference}$ = 98.4%, d = -0.59), we do not interpret this difference as meaningful given its small size. Differences in the slopes between PD OFF and controls were not reliable (-0.01 [-0.02 0.002], $p_{difference}$ = 93.8%, d = -0.44). Therefore, performance of initial forces when the feedback was visible was similar across groups and conditions, reiterating the finding presented in the main results that people with PD are able to produce all of the requested goal forces with their more affected arm.

Next, we analyzed the match forces (Fig. S2b) in the same manner as in the main results: we regressed the mean-centered match force data against the mean-centered initial force data for each participant. However, unlike the main results, when participants with PD used the less impaired arm to match forces (Fig. S2d), we found no differences between groups or conditions for both the intercept (PD ON vs control = -2.7% max force [-12.3 7.3], $p_{difference}$ = 70.4%, d = -0.13; PD OFF vs control = 0.5% max force [-9.9 11.1], $p_{difference}$ = 53.0%, d = 0.02; PD ON vs PD OFF = 0.9% max force [-5.9 7.5], $p_{difference}$ = 61.6%, $d_z$ = 0.06), and the slope (PD ON vs control = -0.04 [-0.18 0.09], $p_{difference}$ = 73.5%, d = -0.16; PD OFF vs control = -0.03 [-0.19 0.13], $p_{difference}$ = 63.6%, d = -0.09; PD ON vs PD OFF = 0.04 [-0.09 0.16], $p_{difference}$ = 70.5%, $d_z$ = 0.10). Thus, force matching with the less affected arm yielded similar performance for both PD and control groups, highlighting that the observed differences in the same-arm force matching task likely arose due to a misperception of effort for the more affected arm during the matching phase of the task, rather than an inability to sense the initial force with the more affected arm.

We also assessed the cognitive perception of effort in the same manner as the first run of the force matching task. Participants rated their perceived effort on a scale of 0 to 10, where 0 represents no effort and 10 represents their maximum possible effort. We analyzed the subjective effort reports by again mean-centering the data, then regressing each participant's mean-centered subjective effort reports against the respective mean-centered initial forces (Fig. S2e). We did not observe reliable differences between PD ON and controls for either the intercept (0.4 [-0.4 1.3], $p_{difference}$ = 85.0%, d = 0.26) or the slope (0.01 [-0.01 0.03], $p_{difference}$ = 84.1%, d = 0.25). We also did not observe differences between the PD ON and PD OFF conditions for either the intercept (-0.2 [-0.8 0.3], $p_{difference}$ = 81.3%, $d_z$ = -0.18) or slope (-0.003 [-0.01 0.01], $p_{difference}$ = 67.2%, $d_z$ = -0.09). However, like the same arm version of the force matching task, we observed greater intercepts for the PD OFF condition compared to controls (0.8 [-0.1 1.7], $p_{difference}$ = 95.5%, d = 0.42). While the slopes were also greater for the PD OFF group compared to controls, this difference was not reliable (0.01 [-0.004 0.03], $p_{difference}$ = 93.3%, d = 0.37). This suggests that the combined effects of PD and medication status affected the cognitive perception of effort even for the less affected arm.

In summary, the sensorimotor perception of effort was disrupted for people with PD only on the more affected arm and was not dopamine dependent, whereas the cognitive perception of effort generalized across both the affected and less affected arms.

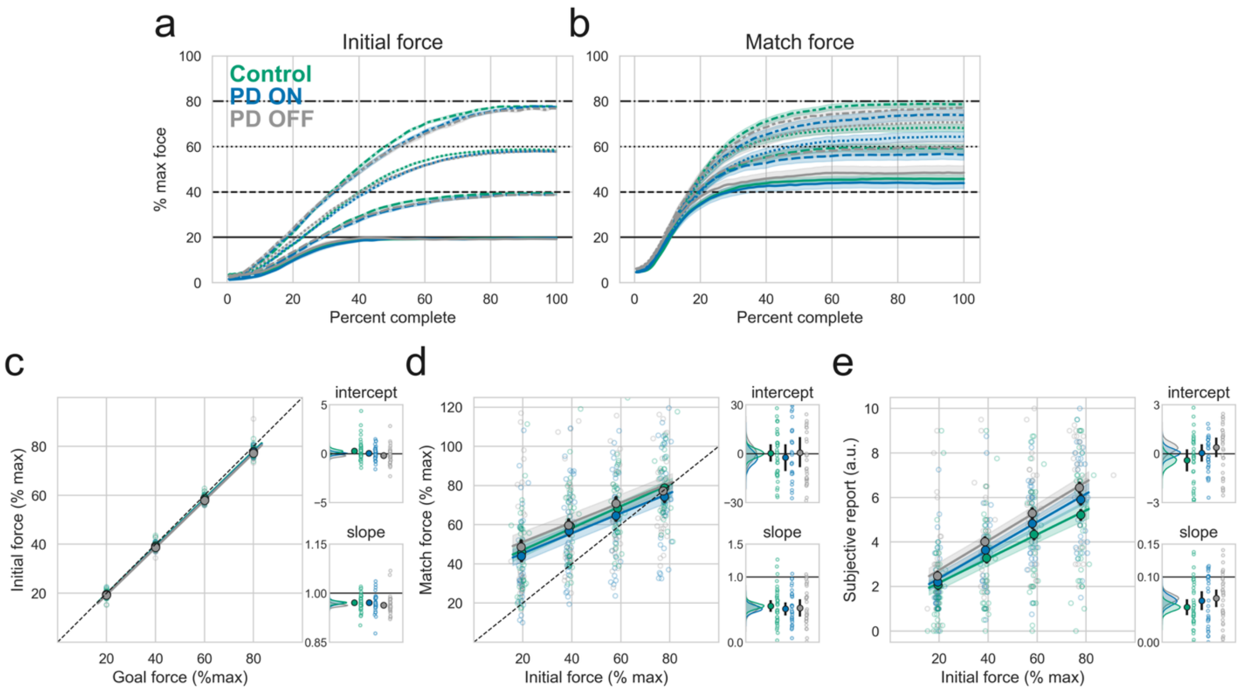


**Figure S2: Perception of effort with the less impaired arm.** Mean initial **(a)** and match **(b)** force traces for each group (colors) and each goal force, denoted with different line styles. For the purposes of these plots, each participant's force trace data was averaged at each percentile so everyone would have 100 data points. Shading represents standard error of the mean, horizontal black lines represent each goal force, with a different style line for each goal. **(c-e, main panels)** Each outcome of interest (y-axis) plotted against the goal or initial forces. Solid circles with black outlines represent the overall group means, error bars (not visible for all datapoints) represent 1 standard error of the mean (SEM), and open circles represent individual participant means at each goal force level. The solid lines are the mean regression lines for each group and shading represents 1 SEM. **(c-e, inset panels)** Kernel density estimates of the posterior distribution of group means for the intercepts (top) and slopes (bottom), calculated using Bayesian t-tests. The empirical group means (filled circles) and individual datapoints (open circles) are plotted to the right. Error bars for these plots (not visible for all datapoints) reflect the 95% HDIs for each respective posterior distribution. Note that the intercepts are calculated from the mean-centered data. **(c)** Performance on the initial forces, when the feedback was visible, was similar across groups and conditions. **(d)** Match forces were similar for all groups and conditions. **(e)** Subjective effort ratings were higher for the PD OFF group compared to controls.
